# Developmental arcs of plasticity in whole movement repertoires of a clonal fish

**DOI:** 10.1101/2023.12.07.570540

**Authors:** Sean M. Ehlman, Ulrike Scherer, David Bierbach, Luka Stärk, Marvin Beese, Max Wolf

## Abstract

Developmental plasticity at the behavioral repertoire level allows animals to incrementally adjust their behavioral phenotypes to match their environments through ontogeny, serving as a lynchpin between ecological factors that cue phenotypic adjustments and evolutionary forces that select upon emergent phenotypic variation. Quantifying the continuous arcs of plasticity throughout animals’ development, however, has often been prohibitively challenging. Here, we leverage recent advancements in high-resolution behavioral tracking and analysis to (i) track the behavior of 45 genetically identical fish clones (*Poecilia formosa*) reared in near-identical environments during their first four weeks of life at 0.2 s resolution and (ii) quantify the continuous arcs of plasticity across entire behavioral repertoires through development. Doing so, we are able to test one of the most fundamental theoretical predictions from Bayesian models of development that in stable (but initially unknown) environments, behavioral plasticity should gradually decrease from a maximum at the beginning of life. Using two approaches to measure plasticity across ontogeny, we first quantify plasticity in individual behavioral metrics before also developing a novel whole-repertoire approach that calculates plasticity as the degree of ‘behavioral entropy’ across a multi-dimensional behavioral phenotype space. We robustly find – despite experimentally matching as best as possible the assumptions of models that predict decreasing plasticity – a ∼two-week initial increase in plasticity in movement behaviors before plasticity subsequently decreased. Our results challenge a common intuition about the optimal developmental course of plasticity through early ontogeny, thereby also demonstrating the value of long-term behavioral tracking approaches for testing fundamental predictions on phenotypic development.

**Significance statement:** Behavioral plasticity across development may help animals adjust to uncertainty in moderately unpredictable environments. In stable environments, developing animals should gradually decrease this uncertainty through ontogeny, becoming less sensitive to incoming information (and thus less behaviorally plastic) as they age. This intuitive expectation of ‘old dog’ inflexibility to ‘new tricks’, however, has not been adequately tested with the long-term, highresolution datasets that would be ideal. Here, we achieve such a test and emphasize the significance of this study in (1) providing a novel method for quantifying multi-dimensional behavioral plasticity continuously across long-term, high-resolution behavioral timeseries and in (2) testing fundamental theory that links the temporal patterning of environmental conditions to evolved patterns of behavioral plasticity across development.

## Introduction

Exploration is key aspect of early life for most animals, providing them with critical information about themselves and their environments. While new information encountered during exploration spurs phenotypic adjustments throughout life, the highest degree of plasticity (or behavioral flexibility) is often expected to occur early in life (1, 2). This early life developmental plasticity may thus potentiate greater phenotypic integration, specialization, and/or competitive advantages compared to conspecifics that delay development (3–6). As development progresses, plasticity’s benefits (e.g., effective exploration of an environment, behavioral pattern, etc.) may be rebalanced against its costs (e.g., increased error rate, opportunity costs, etc.): in moderately stable and predictable environments, exploring environmental cues may not provide much new information (1, 7); aging animals may not have enough time left to capitalize on major plastic adjustments (8); and while plasticity may offer efficient ways to explore an environment or a set of behaviors, this exploring may preclude exploiting already acquired information or further developing existing behavioral patterns (9, 10). This rebalancing of the costs and benefits of plasticity across development governs individuals’ trajectories of plasticity across development and is a fundamental tenet of a large and influential theoretical literature on developmental plasticity (1, 4, 7, 11–13); in sharp contrast, studies that empirically document the arc of developmental plasticity of any particular behavior—let alone multi-dimensional behavioral repertoires—throughout ontogeny have been exceedingly few (due in part to the time- and dataintensive nature of measuring behavior continually through long periods of development).

Our study builds on novel technological approaches to measuring behavior represented by automated tracking in high spatiotemporal resolution, allowing for near-continuous, long-term behavioral observation, and the quantification of multi-dimensional movement repertoires (14– 20). These advances present a path forward towards quantifying continuous plastic changes through development (21) and may help to address a relative dearth of development-length behavioral datasets needed to address theory in this area (22). This study focuses specifically on testing a basic and robust theoretical prediction that, in stable environments, given the potential benefits of exploring environments or behaviors early, plasticity should be highest at the beginning of life and, as ontogeny progresses, should monotonically decrease (7, 13). This prediction relies on the assumption that as animals develop, they improve their ‘estimate’ of the environment and reduce uncertainty through the steady integration of new information that they repeatedly sample, leading to a decrease in sensitivity to new cues. While some empirical tests exist that test these basic predictions (23–26), few take a long-term developmental approach and measure behavior either continuously enough or at a resolution high enough to capture the trajectory of plasticity through ontogeny.

In this paper, we address this gap. Tracking genetically identical fish at 0.2-second resolution from birth until four weeks of life (i.e., ∼1/3 of their total development to sexual maturity) in stable, highly standardized conditions, we develop a novel measure for mapping the continuous arc of behavioral plasticity across development. Whereas behavioral plasticity is typically measured in single behavioral dimensions (e.g., activity, exploration, etc.), we here quantify behavioral plasticity using a whole-repertoire approach that measures plasticity as the degree of Shannon entropy in a high-dimensional behavioral hypervolume. With this wholemovement-repertoire approach for measuring behavioral plasticity across ontogeny, we robustly find that individuals exhibit an inverted U-shape pattern of behavioral plasticity through ontogeny, initially increasing in plasticity until an average of ∼17 days after birth, before then decreasing in behavioral plasticity.

### Behavioral plasticity in individual behavioral metrics through ontogeny

We filmed 45 genetically identical Amazon mollies (*Poecilia formosa*) reared individually in near-identical environments (27) from above, eight hours a day from their first to 28^th^ day of life at five frames per second, yielding a timeseries of approximately 180 million x-y coordinate datapoints of fish in space. From these coordinates, we obtained three timeseries of the most basic behavioral metrics for which the original 0.2-second temporal resolution could be maintained: (1) an instantaneous measure of a fish’s activity, for which we used the Euclidean distance (‘step length’) between two consecutive x-y coordinate points, (2) a measure of a fish’s bearing relative to their previous movement in the preceding timepoint (i.e., their ‘turning angle’), and (3) a measure of a fish’s position in space relative to a salient aspect of their environment, for which we chose the distance to the nearest tank wall.

To quantify an individual’s behavioral plasticity in a particular trait through development, we calculated the coefficient of variation (CoV) for a given trait of a given individual by dividing their trait’s standard deviation by its mean for a given interval of time. Since the means and variances of all three basic behavioral metrics were positively correlated with each other, calculating the CoV for each trait isolated the effect of a trait’s variance from the correlated effect of its mean; this measure, when used to compare changes within a particular trait through time or contexts (rather than among traits), can be used as a measure of phenotypic flexibility or plasticity (28). In order to map the change in CoVs of each of the three basic behavioral metrics over the course of the first 28 days of individuals’ lives, CoVs were calculated over hour-long intervals throughout development for each individual, such that each individual had 8 hours * 28 days = 224 CoV measures for each of the behavioral metrics spanning the first 28 days of life.

Each of an individual’s 224 CoV measures for each metric was thus ultimately based on 18k raw data points (each hour contained 18k raw data points). This yielded a timeseries of behavioral plasticity in each metric spanning the first 28 days of development. Selecting between linear and quadratic mixed models (*Supplemental Tables 2*.*1-3*) using their Akaike and Bayesian Information Criteria revealed that, in all three cases (step length, turning angle, and distance to the tank wall CoVs), quadratic models with inverted U-shapes and a peak of CoV between 110 and 180 hours (between day ∼14-22) of observation (depending on the metric) had the highest support (*Supplemental Table 2*.*4*). This indicated that behavioral plasticity in the three most basic behavioral metrics with time resolution that matches raw x-y coordinate measurements (5 Hz) exhibited an initial increase in behavioral plasticity lasting approximately 2.5 weeks, followed by a subsequent decrease (Fig 1).

**Figure 1.**
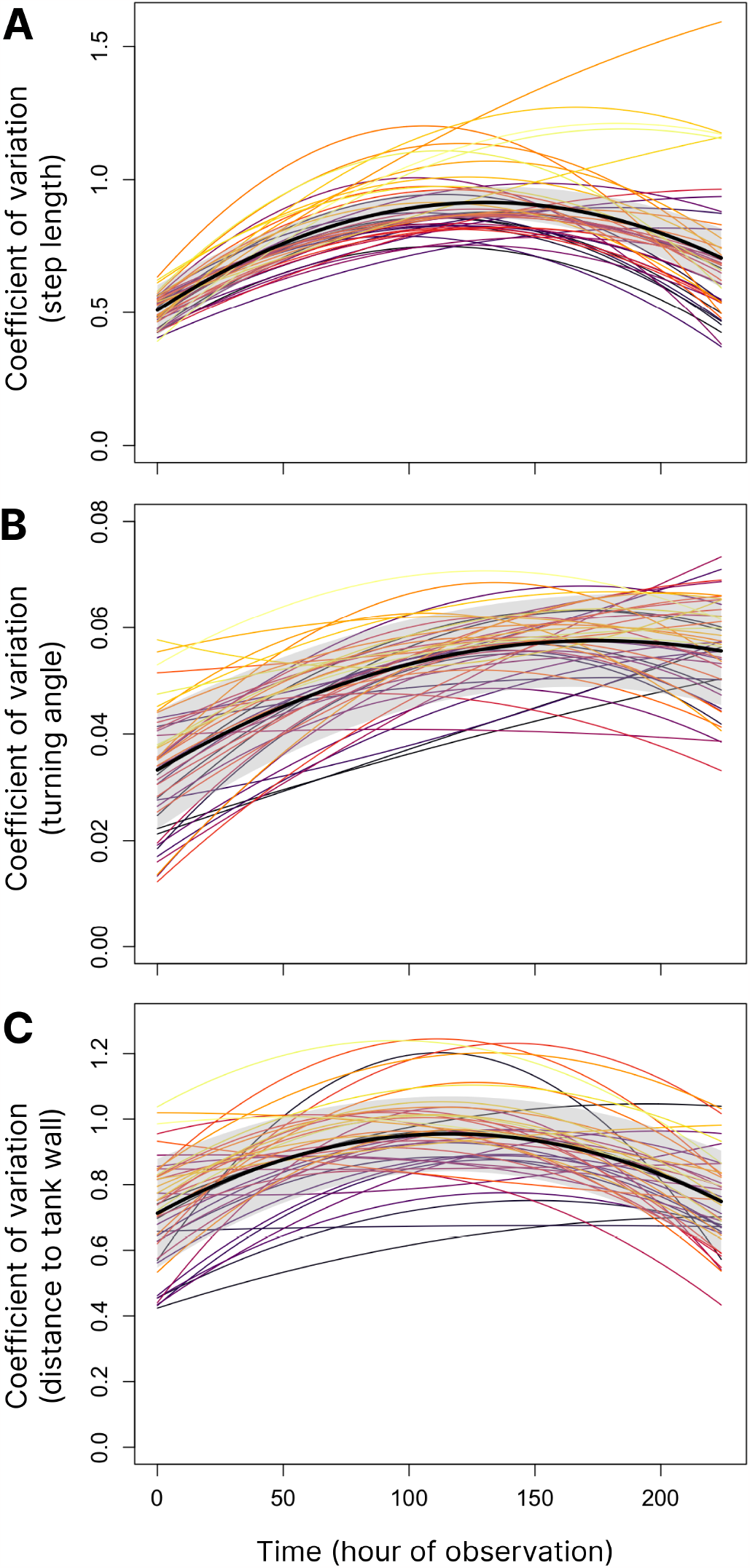
The relationship between developmental time and a measure of behavioral flexibility—the coefficient of variation (CoV)—for three basic measures of behavior: (**A**) a measure of activity (here, ‘step length’), (**B**) a measure of a fish’s bearing (here, ‘turning angle’), and (**C**) a measure of relative spatial position (here, ‘distance to the tank wall’). A 95% confidence interval around the quadratic regression is shown in gray. Colored lines represent individual developmental trajectories obtained from a generalized linear mixed model in which varying intercepts, linear, and quadratic coefficients varied by individual.

### Behavioral plasticity as entropy in multi-dimensional behavioral phenotype space

Our second approach to quantifying trajectories of plasticity across development sought to leverage the high dimensionality in our dataset to represent behavioral plasticity as the diversity of ‘movement’ in multi-dimensional behavioral phenotype space, integrating across the basic behavioral metrics and the multiple scales of temporal autocorrelation potentially present in their timeseries (Fig 2). While the developmental arcs of behavioral plasticity represented by the relationships among developmental time and the CoVs of each of the three basic behavioral metrics provided an important validation step and foundation for further analyses, the multidimensional repertoire approach provided some distinct additional benefits: (i) including all metrics in a single measure of repertoire-wide plasticity allowed for the proportional representation of each metric in their contribution to the overall degree of total plasticity, (ii) while all metrics were represented at 5Hz resolution, plasticity in movement repertoires likely exists across a range of timescales, some much longer; the multi-dimensional repertoire approach thus involved wavelet transforms applied to the timeseries of basic behavioral metrics (29), creating additional feature dimensions that captured the degree of temporal autocorrelation in behavior (which may also represent important axes of behavioral variation, themselves (30)) across longer timescales in the basic behavior metrics (Fig 2B-D; see *Materials and methods*).

**Figure 2.**
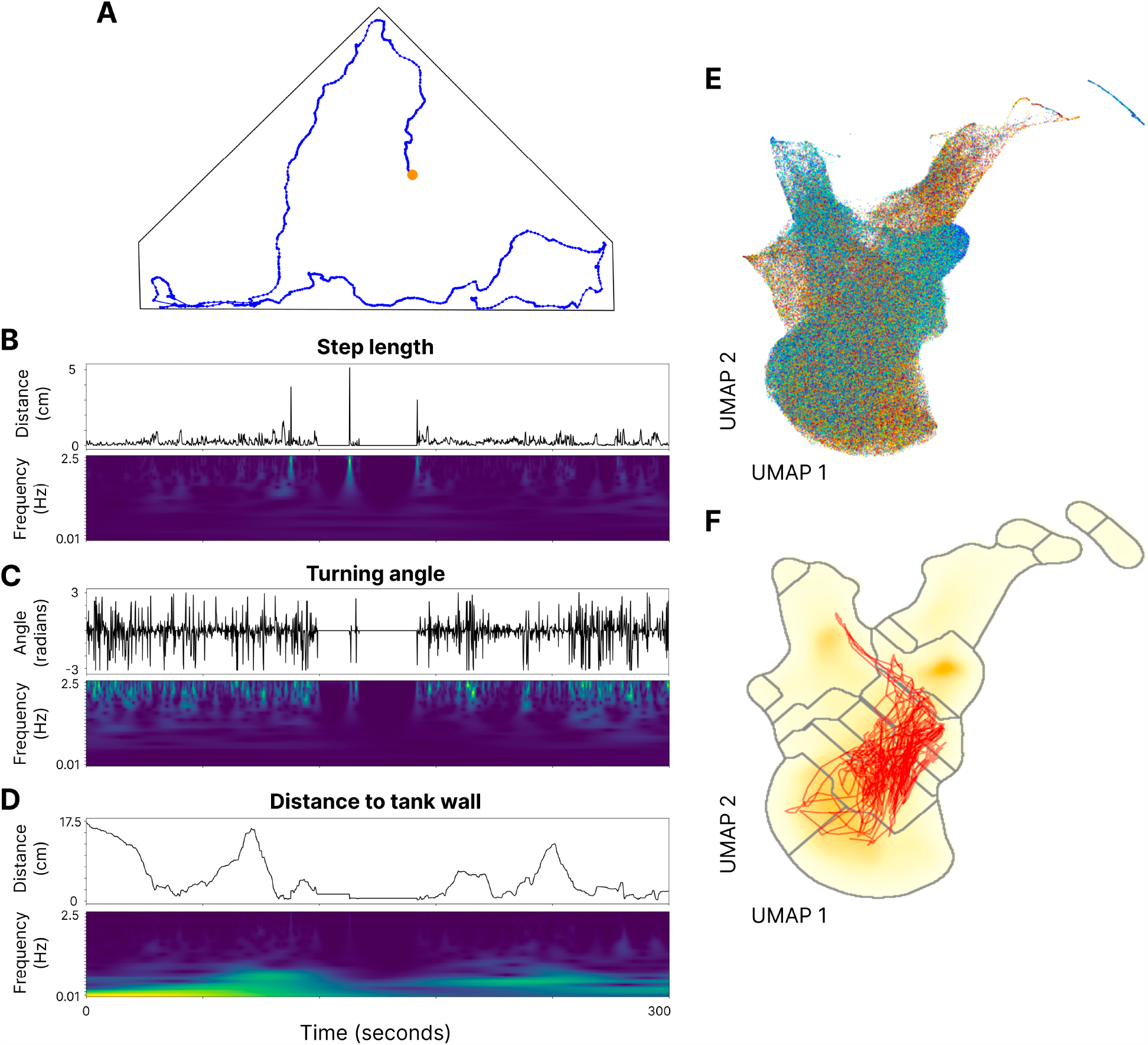
A schematic demonstrating the conversion of an individual’s movement pattern in 2D physical space into a timeseries of ‘movement’ through behavioral phenotype space. (**A**) Here, a 300-second period of an individual’s movement in physical tank space (i.e., triangular shape) is recorded at 0.2-second resolution and plotted as a series of x-y coordinate points. The beginning of the movement sequence is indicated by an orange dot. Step length (**B**), turning angle (**C**), and distance to the nearest tank wall (**D**) are then directly calculated from the raw x-y coordinate data, producing timeseries of these three metrics (upper graph in B-D). Wavelet transforms are then applied to these (normalized) timeseries, with 25 distinct wavelets of frequencies ranging from 0.01-2.5 Hz yielding a 75-dimension feature space. Shown in the lower graph of B-D are spectrograms showing the amplitudes of the 25 Morlet wavelets on each timeseries (minimum amplitudes in dark blue for all spectrograms are 0 s; maximum amplitudes are in bright yellow and correspond to 0.65, 0.40, and 0.16 s for B, C, and D, respectively). (**E**) Data for all time points and all individuals represent a total of ∼180 million measured behavioral datapoints, each represented in a 75-dimensional behavioral phenotype space. Here, the behavioral phenotype space is represented in a dimensionally reduced UMAP embedding where different colors represent the 45 distinct individuals. This UMAP projection is downsampled 500x, and each point has 50% transparency. (**F**) Watershed clustering on the inverse of data density in the UMAP embedding serves as a method to grid behavioral space into *k* (here, 20) grid cells, over which behavioral entropy is calculated. The red trajectory showing ‘movement’ in behavioral phenotype space for a single individual corresponds to the 300-second timeseries depicted in A-D.

After wavelet transforms were performed, data from all individuals and all timepoints throughout development represented a 75-dimensional behavioral phenotype space (Fig 2E). In order to quantify behavioral plasticity in this phenotype space over time, we calculated a measure of the spread of behaviors exhibited by any particular individual as their Shannon ‘behavioral entropy’. This same basic entropy measure (31) is commonly used in biology to calculate biodiversity across a species assemblage (32) or heterogeneity of physical space use in a landscape (33, 34), and has also been applied to measure behavioral variability among a range of pre-defined behaviors (35) and molecular diversity across genomes (36); here we extend the application of entropy as a measure of an individual’s behavioral plasticity when entropy is calculated across a multi-dimensional, cluster-delineated behavioral phenotype space as a continuous function of time. Since Shannon entropy calculations in space generally require ‘space’ to be discretized as a grid, gridding behavioral phenotype space was performed using unsupervised behavioral clustering via UMAP (Uniform Manifold Approximation and Projection (37)) embedding, Gaussian smoothing of data density in embedding space, and a watershed transform on the inverse of that density space (38)(Fig 2F). Entropy was calculated over hourlong intervals through development for each individual, as with CoV measures, yielding 224 measures of entropy spanning the first 28 days of life for each individual that were each represented in 75-dimensional behavior space.

As with CoV measures, the developmental time course of entropy was best described as an inverted U-shape (*Supplementary Tables 2*.*3-4*): Figure 3A shows developmental arcs of behavioral entropy in which behavioral plasticity initially increases, peaking around hour 135 (∼17 days) before then decreasing. Individual fish (colored lines) vary in their mean entropy through development, and it is instructive contrasting behavioral patterns of relatively low- and high-entropy individuals. Fig 3B shows, for example, that individuals with low behavioral entropy exhibit patterns of reduced ‘movement’ in behavioral space, with some low entropy individuals spending up to 90% of their time in any given hour in a single behavioral cluster (Fig 3B). High entropy individuals, in contrast, exhibit a more uniform distribution of cluster occupancy, spending approximately equal amounts of time in a larger number of behavioral clusters during each hour. This can be represented as trajectories through behavioral phenotype space (Fig 3C): even though any two low entropy individuals might exhibit quite different individual behaviors (i.e., be segregated in behavioral phenotype space), they are united in their limited movement through total behavioral space, in contrast to high entropy individuals who use a much greater proportion of behavioral phenotype space. The patterns of greater ‘movement’ in behavioral space (and thus higher behavioral entropy) correspond to a greater diversity of movement patterns in physical space when simultaneous fish movements in physical space and behavioral (UMAP) space are visualized side-by-side (*Supplementary Movie 5*.*1*). These results are further validated by and extend the standard but more limited approach to quantifying patterns of single-behavioral-dimension plasticity (CoVs) in the basic behavioral metrics shown earlier (Fig 1).

**Figure 3.**
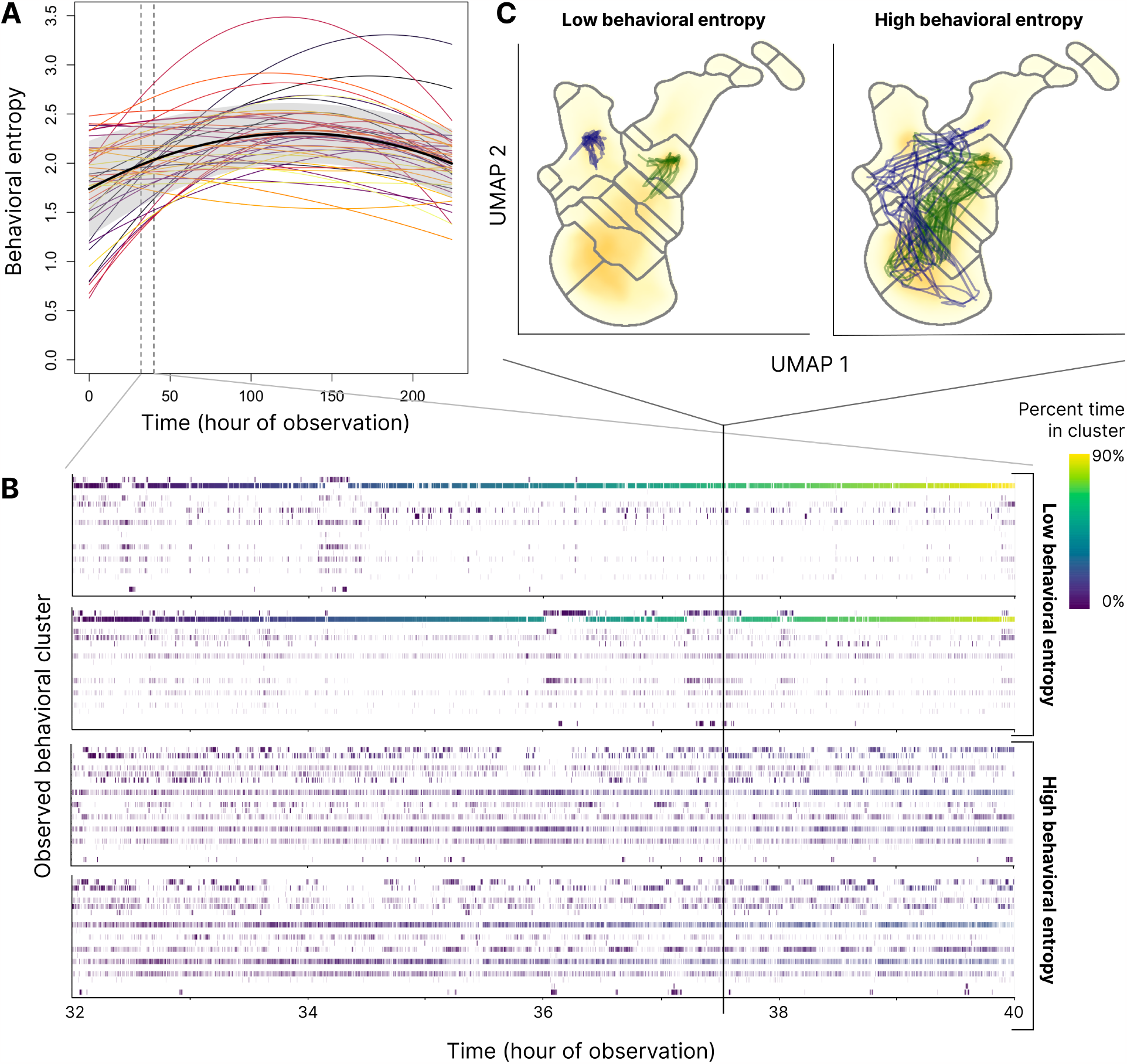
The developmental arc of plasticity across the first 28 days of life for individuals as represented by Shannon entropy calculated across a behavioral landscape. (**A**) Average behavioral entropy in the population initially increases, peaking around observation hour 135 (∼17 days after birth), before then decreasing. Colored lines represent individual developmental trajectories obtained from a generalized linear mixed model in which varying intercepts, linear, and quadratic coefficients varied by individual. (**B**) Representative day-length ethograms (from 8 observation hours of day 5) in which cluster occupancy (cluster 1-20) is indicated through time at 0.2-second resolution by small vertical ticks. The top two ethograms demonstrate a pattern of low behavioral entropy over this interval, while the bottom two ethograms represent a pattern of high entropy. (**C**) Three-minute trajectories (corresponding in time to the vertical solid line shown in (B)) through behavioral phenotype space represented as a non-linear embedding via UMAP. Watershed cluster boundaries are shown; the two low entropy individuals shown in (B) are represented in the left UMAP, while the high entropy individuals shown in (B) are in the right UMAP.

## Discussion

We mapped the developmental arcs of behavioral plasticity in entire movement repertoires of 45 genetically identical fish reared in near-identical environments for the first 28 days of their lives at 0.2 s resolution. Employing a novel measure of entropy in behavioral repertoire space, we then tested a key prediction from Bayesian models of development that, in stable environments behavioral plasticity should gradually decrease from a maximum at the beginning of life (7, 13, 39). In our experiment setup, fish experienced constant and benign conditions from their day of birth through the duration of the experiment; despite this matching as best as possible the assumptions of models that predict decreasing plasticity, we found a ∼two-week initial increase in plasticity in movement behaviors before plasticity subsequently decreased.

This deviation of our empirical results from those expected by theory might be explained by discontinuities between informational priors (e.g., set evolutionarily or via parental effects) and individually sampled information (40–44). It is possible, for example, that newborn individuals — small prey fish vulnerable to high levels of predation in natural streams (45) — start life with a relatively conservative ‘prior’ about the level of risk, and thus show an initial increase in plasticity early in life as they sample to rebalance their erroneous estimate of risk levels in now-predator-free, benign lab conditions. Alternatively, while predicted developmental trajectories of plasticity in Bayesian models of development are driven primarily by informational constraints (e.g., trade-offs involved in ‘exploring’ new information or ‘exploiting’ already sampled information), there may be other developmental constraints, such as motor or cognitive constraints (46, 47), that otherwise restrict the behavioral repertoire range of newly born mollies, delaying periods of maximal plasticity till later in development (48). Whatever the underlying cause explaining the deviation of our empirical results from those predicted by basic Bayesian theory, we emphasize the utility in applying recent advancements in high-resolution behavioral tracking to test and refine open questions in the theory of behavioral development.

Our results add to and significantly extend a nascent but growing set of studies using high-throughput behavior tracking and multi-dimensional behavioral representations to document behavioral repertoire changes throughout time (49–55); in particular, our results demonstrate the value of applying these novel technologies to the continuous measurement of behavioral plasticity during ontogeny. This is particularly important given the long-held consensus that more continuous measurements of behavior and its plasticity through development are critical in advancing our understanding of the ecological and evolutionary factors shaping behavior and its development (22, 56). Our results for quantifying and analyzing developmental arcs of behavioral plasticity could be productively applied to a range of important questions in behavioral development, including: the development of individual differences in plasticity (57–60), developmental constraints on patterns of behavioral variation (61), and the role of developmental plasticity in mediating behavioral responses to environmental change (62– 64). Given the crucial role of developmental plasticity in both translating ecological cues into phenotypic effects and producing the range of phenotypes upon which selection and evolution act (65–67), new tools for fully mapping the continuous arcs of such developmental plasticity in behavior are likely to significantly contribute to our overall understanding of the ecological and evolutionary causes and consequences of behavioral development.

## Materials and methods

### Study species and experimental design

All animal care and experimental protocols complied with local and federal laws and guidelines and were approved by the appropriate governing body in Berlin, Germany, the Landesamt fur Gesundheit und Soziales (LaGeSo G-0224/20).

Details on study species, animal housing and feeding protocols, and experimental design are described in (27); we briefly describe them here. In this study, genetically identical gravid Amazon mollies (*Poecilia formosa*) were isolated from a single isogenic stock kept at Humboldt Universität zu Berlin (Berlin, Germany). The Amazon molly is a gynogenetic species of freshwater fish (68, 69) — the first described species of clonal vertebrate (70) – with a diverse behavioral repertoire through development (27, 71, 72). Offspring from three of these isogenic mothers were used as experimental animals in behavioral observations. In addition to the original three mothers of experimental animals being genetically identical and arising from the same stock tank, we also accounted for individual mother ID in all statistical models (*Supplements 2-4*). In this way, we both minimized any maternal effects due to obvious differential experiences of mothers and ensured that all experimental animals were genetically identical. In total, three mothers provided 45 experimental fish, which were transferred on the day of their birth to large individual observation tanks (*Supplementary Fig 1*.*1*). From the next day (their first full day of life) until an age of 28 days, fish were filmed from above using Basler acA5472 cameras at five frames per second continuously for eight hours per day. Given that mollies reach sexual maturity at approximately three months, behavioral observations covered a full third of development to maturity for these fish. Note that while overhead filming meant that the third dimension of movement (up and down in the water column) was not recorded, the water level in tanks was kept relatively shallow (at a depth of ∼7cm) so that the dominant dimensions of possible movement were largely confined to two dimensions. Fish were kept on a 12:12h light:dark cycle with an air temperature of approximately 24 ± 1°C and fed daily during a two hour period following the 8-hour filming period with a stationary ‘food patch’ consisting of Sera vipan baby fish food fixed in agar. The white polyethylene observation tanks were illuminated from below, allowing for ease of automated video tracking conducted using the software Biotracker (73). This generated data in the form of a timeseries of fish positions in x-y coordinate space with 0.2 second temporal resolution.

### Statistical models for basic behavioral measures

For each basic behavioral metric (i.e., step length, turning angle, and distance to the tank wall), we calculated a trait’s coefficient of variation (CoV) by dividing its standard deviation by its mean for a given interval of time at hour-length intervals (constituting 18k raw data points for each CoV datapoint). See *Supplement 3* for further analyses using CoVs calculated over daylength intervals; note that results were robust to the specific interval used to calculate individual CoV datapoints). Visual inspection of the course of plasticity in all three basic behavioral traits revealed inverted U-shapes rather than monotonic decreases in plasticity; this was confirmed by model comparison between linear and quadratic models in a mixed modeling framework with a main fixed effect of developmental time, controlling for mother ID, tank position within the experimental room (center or periphery within a flow-through system), and tank system (one of four flow-through systems) as additional fixed effects. Individual ID was treated as a random effect, with randomly varying intercepts and slopes in the case of linear models and randomly varying intercepts, linear, and quadratic coefficients in the case of quadratic models (see *Supplement 2* for further details).

### Multi-dimensional behavioral phenotype space

In order to create a multi-dimensional behavioral phenotype space, we first performed wavelet transforms on each of the three (normalized) basic behavioral metric timeseries following (29, 74): 25 wavelets of different frequencies (min. frequency: 0.01 Hz; max. frequency: 2.5 Hz) were convolved over the three basic behavioral measure timeseries yielding a total of 75 feature dimensions. The convolution of a wavelet over a behavioral timeseries yields an ‘activation’ at each point in the timeseries, which is a measure of the waveform’s ‘goodness of fit’ between a timeseries and that waveform centered at a particular point along that timeseries (75). The timeseries of a wavelet’s activations filtered over a behavioral timeseries thus becomes its own timeseries. By using a basic waveform with short-lived localized oscillations (here, the Morlet wavelet (76)), changing the frequency of that waveform, and comparing activations of these waveforms with different frequencies on a behavioral timeseries, variable patterns of temporal autocorrelation in the data can be captured as additional features in behavioral repertoire space. Thus, by wavelet transforming spatial positioning, activity, and turning angles, we can jointly represent behavioral phenotype space as encompassing variation in the data of where animals are, how and how quickly they move, as well as the varying degrees of temporal autocorrelation in these measures.

### Behavioral entropy measure

After wavelet transforms were performed, data from all individuals and all timepoints through development represented a 75-dimensional behavioral phenotype space. In order to quantify ‘movement’ through this phenotype space over time, we calculated a measure of the spread of behaviors exhibited by any particular individual over all of behavioral space as their ‘behavioral entropy’: the greater behavioral diversity individuals exhibited across the total behavioral space, the greater their behavioral entropy and thus plasticity. In order to calculate entropy, behavioral space first needed to be partitioned, which was performed through the use of a non-linear dimension reduction step (Uniform Manifold Approximation and Projection; UMAP (37)) followed by Gaussian smoothing of data density in embedding space and a watershed transform on the inverse of that density space (38). This yielded a specified number of behavioral clusters; for the main analysis, we used 20 clusters (i.e., 20 ‘grid cells’ over which entropy was calculated), but results were robust to much lower cluster numbers (see *Supplement 4*).

*Supplement 4* also contains additional analyses with an alternative clustering algorithm (k-means performed over the full 75-dimension behavioral space), and results were again qualitatively unchanged.

Once behavioral clusters were identified, Shannon entropy over behavioral space, *H*(*X*), was calculated as:

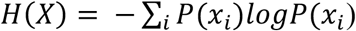

where *P*(*x*_*i*_) is the probability of being in any cluster, *i*, computed from the data as the proportion of time a fish ‘visited’ a particular cluster (i.e., exhibited a behavior within a behavioral cluster). In order to calculate the timeseries of entropies for each individual fish—and thus plasticity in their movement repertoire—over development, hourly entropy for each fish was calculated by binning behavioral timeseries into one-hour intervals. Note that behavioral entropy was also calculated on a day-length interval (*Supplement 3*); results do not differ qualitatively across these interval scales. Lastly, in order to assess the specific shape of developmental arcs in behavioral plasticity (i.e., entropy), we again employed a model comparison approach between linear and quadratic models in a mixed modeling framework with a main fixed effect of developmental time, controlling for mother ID, tank position within the experimental room (center or periphery within a flow-through system), and tank system (one of four flow-through systems) as additional fixed effects. Individual ID was treated as a random effect, with randomly varying intercepts and slopes in the case of linear models and randomly varying intercepts, linear, and quadratic coefficients in the case of quadratic models (see *Supplement 2* for further details). All linear mixed effects models (for both CoV measures and entropy) were conducted using the lme4 package (77) in R version 4.1.1 (78). All other analyses (e.g., wavelet transforms, UMAP embedding and watershed clustering) were performed in Python version 3.11.0.

## Acknowledgments

We gratefully acknowledge funding from the Deutsche Forschungsgemeinschaft under Germany’s Excellence Strategy EXC 2002/1, ‘Science of Intelligence’ (project number #390523135). Members of the Wolf and Krause labs at the Leibniz Institute of Freshwater Ecology and Inland Fisheries, Humboldt University, and SCIoI Excellenzcluster provided invaluable help with animal husbandry, experimental design and logistics, as well as coding and data management: particular thanks are extended to F. Francisco, J. Krause, C. Schutz, D. Strasiewsky, J. Piotrowski, O. O’Connor, and R. Leipold. We are grateful also to N. Walasek for helpful discussions and comments on the manuscript.

## Notes

**Funding:** This work was supported by the Deutsche Forschungsgemeinschaft (DFG) under Germany’s Excellence Strategy – EXC 2002/1 ‘Science of Intelligence’ – project number 390523135.

### Competing Interest Statement

The authors have declared no competing interest.

