## Supplementary Material for "Developmental arcs of plasticity in whole movement repertoires of a clonal fish"

for

**CONTENTS**

Supplement 1: Observation tank setup ……………………………………………………...2

Supplement 2: Model output and selection……………………………………………………..3

Supplement 3: Developmental arcs of plasticity with day-length time intervals…………………7

Supplement 4: Behavioral entropy measures calculated across varying cluster numbers and with multiple clustering algorithms……………………………………………………….11

Supplement 5: Movie of simultaneous movement in physical space and behavioral phenotype space for low- and high-entropy individuals………………..…………………15

### Supplement 1: Observation tank setup

**Supplementary Figure 1.1.** The layout of the experimental tanks in which fish were housed and filmed from above for the first 28 days of life. Each experimental unit (shown here) contained two experimental tanks, and experimental units were connected via flow-through water systems. Food patches were only present during the two-hour feeding period, outside of the eight-hour behavioral observation periods included in analyses. Reproduced with permission from (27).

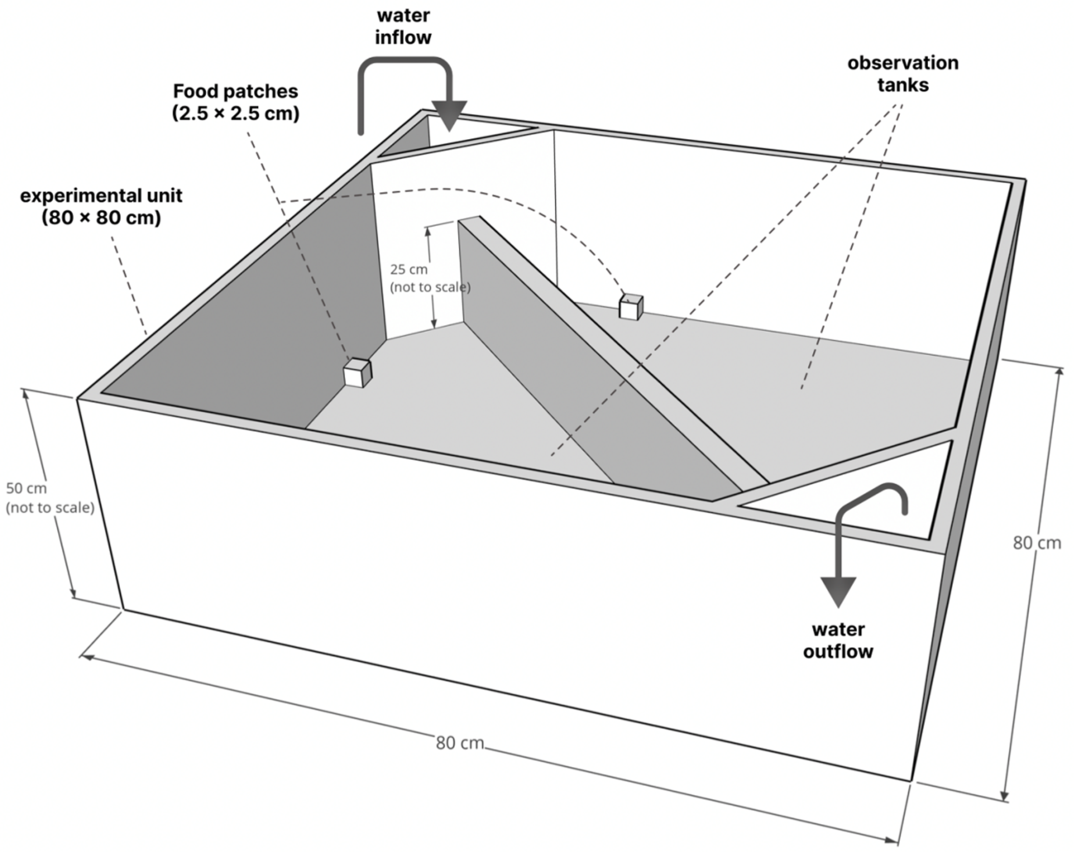

### Supplement 2: Model output and selection

**Supplementary Table 2.1.** Output for the linear models of the coefficient of variation (CoV) for step length, turning angle, and distance to the tank wall regressed against time (in hours) as the continuous fixed predictor of interest. Mother ID, tank position, and tank system are included as categorical fixed predictors, and fish ID is included as a random factor with varying slopes and intercepts.

| | **Step length CoV (linear model)** | | | **Turning angle CoV (linear model)** | | | **Distance to tank wall CoV (linear model)** | | |
|---|---|---|---|---|---|---|---|---|---|
| *Predictors* | *Estimates* | *CI* | *p* | *Estimates* | *CI* | *p* | *Estimates* | *CI* | *p* |
| (Intercept) | 0.696 | 0.612 – 0.779 | **<0.001** | 0.040 | 0.034 – 0.045 | **<0.001** | 0.850 | 0.761 – 0.939 | **<0.001** |
| timestep | 0.085 | 0.046 – 0.123 | **<0.001** | 0.010 | 0.008 – 0.012 | **<0.001** | 0.014 | -0.012 – 0.040 | 0.292 |
| mother ID [2] | -0.013 | -0.116 – 0.091 | 0.813 | -0.004 | -0.011 – 0.003 | 0.230 | -0.125 | -0.234 – -0.015 | **0.025** |
| mother ID [3] | -0.051 | -0.105 – 0.004 | 0.069 | 0.004 | 0.001 – 0.008 | **0.014** | -0.007 | -0.064 – 0.051 | 0.819 |
| tank position [middle] | 0.010 | -0.053 – 0.074 | 0.755 | 0.001 | -0.003 – 0.005 | 0.695 | -0.007 | -0.073 – 0.060 | 0.846 |
| tank position [wall] | -0.004 | -0.071 – 0.063 | 0.912 | 0.001 | -0.004 – 0.005 | 0.807 | 0.005 | -0.066 – 0.075 | 0.900 |
| tank system [2] | 0.010 | -0.062 – 0.081 | 0.789 | -0.001 | -0.005 – 0.004 | 0.734 | 0.075 | -0.000 – 0.150 | 0.051 |
| tank system [3] | 0.006 | -0.070 – 0.081 | 0.883 | 0.001 | -0.004 – 0.005 | 0.814 | 0.018 | -0.062 – 0.097 | 0.665 |
| tank system [4] | -0.029 | -0.101 – 0.043 | 0.424 | -0.003 | -0.008 – 0.001 | 0.183 | 0.001 | -0.075 – 0.076 | 0.989 |
| **Random Effects** | | | | | | | | | |
| $\sigma^2$ | 0.17589 | | | 0.00009 | | | 0.05574 | | |
| $\tau_{00}$ | 0.01916 $_{id}$ | | | 0.00014 $_{id}$ | | | 0.02714 $_{id}$ | | |
| $\tau_{11}$ | 0.01550 $_{id.timestep}$ | | | 0.00004 $_{id.timestep}$ | | | 0.00734 $_{id.timestep}$ | | |
| ICC | 0.07270 | | | 0.36940 | | | 0.18616 | | |
| N | 45 $_{id}$ | | | 45 $_{id}$ | | | 45 $_{id}$ | | |
| Observations | 9861 | | | 9861 | | | 9861 | | |
| Marginal $R^2$ / Conditional $R^2$ | 0.020 / 0.091 | | | 0.245 / 0.524 | | | 0.031 / 0.211 | | |

**Supplementary Table 2.2.** Output for the quadratic models of the coefficient of variation (CoV) for step length, turning angle, and distance to the tank wall regressed against time and time$^2$ (in hours) as the continuous fixed predictors of interest. Mother ID, tank position, and tank system are included as categorical fixed predictors, and fish ID is included as a random factor with varying intercepts, quadratic, and linear coefficients.

| | **Step length CoV (quadratic model)** | | | **Turning angle CoV (quadratic model)** | | | **Distance to tank wall CoV (quadratic model)** | | |
|---|---|---|---|---|---|---|---|---|---|
| *Predictors* | *Estimates* | *CI* | *p* | *Estimates* | *CI* | *p* | *Estimates* | *CI* | *p* |
| (Intercept) | 0.509 | 0.414 – 0.605 | **<0.001** | 0.033 | 0.022 – 0.044 | **<0.001** | 0.713 | 0.547 – 0.879 | **<0.001** |
| timestep^2 | -0.238 | -0.276 – -0.200 | **<0.001** | -0.008 | -0.010 – -0.006 | **<0.001** | -0.178 | -0.215 – -0.141 | **<0.001** |
| timestep | 0.621 | 0.543 – 0.700 | **<0.001** | 0.028 | 0.023 – 0.032 | **<0.001** | 0.415 | 0.328 – 0.503 | **<0.001** |
| mother ID [2] | -0.202 | -0.332 – -0.072 | **0.002** | -0.014 | -0.029 – 0.001 | 0.072 | -0.239 | -0.467 – -0.011 | **0.040** |
| mother ID [3] | -0.058 | -0.125 – 0.009 | 0.087 | 0.003 | -0.005 – 0.011 | 0.429 | 0.072 | -0.048 – 0.192 | 0.238 |
| tank position [middle] | 0.007 | -0.071 – 0.085 | 0.869 | 0.001 | -0.008 – 0.010 | 0.819 | 0.003 | -0.137 – 0.142 | 0.972 |
| tank position [wall] | -0.011 | -0.093 – 0.071 | 0.790 | -0.003 | -0.013 – 0.007 | 0.538 | -0.092 | -0.239 – 0.055 | 0.219 |
| tank system [2] | 0.040 | -0.048 – 0.128 | 0.371 | 0.006 | -0.005 – 0.016 | 0.267 | 0.060 | -0.097 – 0.217 | 0.452 |
| tank system [3] | -0.015 | -0.108 – 0.078 | 0.757 | 0.001 | -0.010 – 0.012 | 0.825 | -0.063 | -0.229 – 0.103 | 0.459 |
| tank system [4] | -0.024 | -0.112 – 0.065 | 0.601 | -0.003 | -0.014 – 0.008 | 0.578 | 0.002 | -0.156 – 0.160 | 0.981 |
| **Random Effects** | | | | | | | | | |
| $\sigma^2$ | 0.16549 | | | 0.00008 | | | 0.04964 | | |
| $\tau_{00}$ | 0.00688 $_{id.2}$ | | | 0.00016 $_{id.2}$ | | | 0.03453 $_{id.2}$ | | |
| $\tau_{11}$ | 0.04112 $_{id.timestep}$ | | | 0.00023 $_{id.timestep}$ | | | 0.08048 $_{id.timestep}$ | | |
| | 0.01136 $_{id.I(timestep^2)}$ | | | 0.00004 $_{id.I(timestep^2)}$ | | | 0.01423 $_{id.I(timestep^2)}$ | | |
| ICC | 0.29347 | | | 0.82355 | | | 0.73049 | | |
| N | 45 $_{id}$ | | | 45 $_{id}$ | | | 45 $_{id}$ | | |
| Observations | 9861 | | | 9861 | | | 9861 | | |
| Marginal $R^2$ / Conditional $R^2$ | 0.056 / 0.333 | | | 0.146 / 0.849 | | | 0.072 / 0.750 | | |

**Supplementary Table 2.3.** Output for linear and quadratic models of behavioral entropy (calculated on a grid of 20 clusters obtained via UMAP dimension reduction and watershed segmentation) regressed against time (in hours) as the continuous fixed predictor of interest. The quadratic model contains an additional term for time$^2$ as a continuous fixed predictor. Mother ID, tank position, and tank system are included as categorical fixed predictors, and fish ID is included as a random factor with varying intercepts and linear coefficients (as well as varying quadratic coefficients in the case of the quadratic model).

| | **Behavioral entropy (linear model)** | | | **Behavioral entropy (quadratic model)** | | |
|---|---|---|---|---|---|---|
| *Predictors* | *Estimates* | *CI* | *p* | *Estimates* | *CI* | *p* |
| (Intercept) | 1.68 | 1.54 – 1.82 | **<0.001** | 1.74 | 1.25 – 2.23 | **<0.001** |
| timestep | 0.11 | 0.03 – 0.19 | **0.007** | 0.88 | 0.63 – 1.14 | **<0.001** |
| mother ID [2] | -0.13 | -0.23 – -0.02 | **0.017** | -1.15 | -1.82 – -0.47 | **0.001** |
| mother ID [3] | -0.08 | -0.13 – -0.02 | **0.005** | -0.23 | -0.59 – 0.12 | 0.201 |
| tank position [middle] | 0.02 | -0.05 – 0.08 | 0.593 | -0.12 | -0.53 – 0.29 | 0.570 |
| tank position [wall] | 0.03 | -0.03 – 0.10 | 0.306 | -0.33 | -0.76 – 0.11 | 0.143 |
| tank system [2] | 0.06 | -0.01 – 0.13 | 0.087 | 0.29 | -0.18 – 0.75 | 0.222 |
| tank system [3] | 0.01 | -0.06 – 0.09 | 0.754 | -0.19 | -0.68 – 0.31 | 0.460 |
| tank system [4] | -0.03 | -0.10 – 0.05 | 0.491 | -0.19 | -0.66 – 0.28 | 0.434 |
| timestep^2 | | | | -0.34 | -0.43 – -0.25 | **<0.001** |
| **Random Effects** | | | | | | |
| $\sigma^2$ | 0.12 | | | 0.10 | | |
| $\tau_{00}$ | 0.17 $_{id}$ | | | 0.32 $_{id.2}$ | | |
| $\tau_{11}$ | 0.08 $_{id.timestep}$ | | | 0.74 $_{id.timestep}$ | | |
| | | | | 0.08 $_{id.I(timestep^2)}$ | | |
| ICC | 0.27 | | | 0.93 | | |
| N | 45 $_{id}$ | | | 45 $_{id}$ | | |
| Observations | 9861 | | | 9861 | | |
| Marginal $R^2$ / Conditional $R^2$ | 0.048 / 0.306 | | | 0.097 / 0.934 | | |

**Supplementary Table 2.4.** Table of Akaike Information Criteria (AIC) and Bayesian Information Criteria (BIC) used to compare linear and quadratic models (see Supplemental Tables 2.1 and 2.2) for each response.

| **Model** | **ΔAIC** | **ΔBIC** |
|---|---|---|
| Step length CoV quadratic | 0.00 | 0.00 |
| Step length CoV linear | 468.68 | 461.48 |
| Turning angle CoV quadratic | 0.00 | 0.00 |
| Turning angle CoV linear | 1250.12 | 1242.92 |
| Distance to tank wall CoV quadratic | 0.00 | 0.00 |
| Distance to tank wall CoV linear | 865.40 | 858.20 |
| Behavioral entropy quadratic | 0.00 | 0.00 |
| Behavioral entropy linear | 1801.03 | 1793.83 |

<u>Supplement 3: Developmental arcs of plasticity with day-length time intervals</u>

**Supplementary Table 3.1.** Output for the linear models of the coefficient of variation (CoV) for step length, turning angle, and distance to the tank wall regressed against time (in days) as the continuous fixed predictor of interest. Mother ID, tank position, and tank system are included as categorical fixed predictors, and fish ID is included as a random factor with varying slopes and intercepts.

| | **Step length CoV (linear model – day interval)** | | | **Turning angle CoV (linear model – day interval)** | | | **Distance to tank wall CoV (linear model – day interval)** | | |
|---|---|---|---|---|---|---|---|---|---|
| *Predictors* | *Estimates* | *CI* | *p* | *Estimates* | *CI* | *p* | *Estimates* | *CI* | *p* |
| (Intercept) | 0.762 | 0.673 – 0.851 | **<0.001** | 0.041 | 0.035 – 0.046 | **<0.001** | 0.890 | 0.796 – 0.983 | **<0.001** |
| timestep | 0.556 | 0.218 – 0.894 | **0.001** | 0.077 | 0.061 – 0.092 | **<0.001** | 0.139 | -0.085 – 0.363 | 0.223 |
| mother ID [2] | -0.027 | -0.140 – 0.085 | 0.637 | -0.004 | -0.011 – 0.003 | 0.229 | -0.131 | -0.246 – -0.016 | **0.025** |
| mother ID [3] | -0.013 | -0.072 – 0.045 | 0.656 | 0.004 | 0.001 – 0.008 | **0.018** | -0.013 | -0.073 – 0.048 | 0.684 |
| tank position [middle] | 0.003 | -0.066 – 0.071 | 0.934 | 0.001 | -0.003 – 0.005 | 0.769 | -0.010 | -0.080 – 0.060 | 0.785 |
| tank position [wall] | -0.004 | -0.076 – 0.068 | 0.911 | 0.000 | -0.004 – 0.005 | 0.836 | -0.008 | -0.082 – 0.066 | 0.837 |
| tank system [2] | 0.013 | -0.064 – 0.091 | 0.731 | -0.001 | -0.005 – 0.004 | 0.707 | 0.062 | -0.017 – 0.141 | 0.122 |
| tank system [3] | 0.009 | -0.073 – 0.091 | 0.831 | 0.001 | -0.004 – 0.005 | 0.784 | -0.004 | -0.087 – 0.080 | 0.929 |
| tank system [4] | -0.027 | -0.105 – 0.050 | 0.488 | -0.003 | -0.008 – 0.001 | 0.177 | 0.001 | -0.079 – 0.081 | 0.978 |
| **Random Effects** | | | | | | | | | |
| $\sigma^2$ | 0.04918 | | | 0.00006 | | | 0.02545 | | |
| $\tau_{00}$ | 0.01522 $_{id}$ | | | 0.00013 $_{id}$ | | | 0.02608 $_{id}$ | | |
| $\tau_{11}$ | 1.05240 $_{id.timestep}$ | | | 0.00252 $_{id.timestep}$ | | | 0.43676 $_{id.timestep}$ | | |
| ICC | 0.24630 | | | 0.47398 | | | 0.32466 | | |
| N | 45 $_{id}$ | | | 45 $_{id}$ | | | 45 $_{id}$ | | |
| Observations | 1239 | | | 1239 | | | 1239 | | |
| Marginal $R^2$ / Conditional $R^2$ | 0.033 / 0.272 | | | 0.295 / 0.629 | | | 0.054 / 0.361 | | |

**Supplementary Table 3.2.** Output for the quadratic models of the coefficient of variation (CoV) for step length, turning angle, and distance to the tank wall regressed against time and time$^2$ (in days) as the continuous fixed predictors of interest. Mother ID, tank position, and tank system are included as categorical fixed predictors, and fish ID is included as a random factor with varying intercepts, quadratic, and linear coefficients.

| | **Step length CoV (quadratic model – day interval)** | | | **Turning angle CoV (quadratic model – day interval)** | | | **Distance to tank wall CoV (quadratic model – day interval)** | | |
|---|---|---|---|---|---|---|---|---|---|
| *Predictors* | *Estimates* | *CI* | *p* | *Estimates* | *CI* | *p* | *Estimates* | *CI* | *p* |
| (Intercept) | 0.521 | 0.419 – 0.623 | **<0.001** | 0.035 | 0.026 – 0.045 | **<0.001** | 0.686 | 0.564 – 0.808 | **<0.001** |
| timestep^2 | -14.09 | -16.250 – -11.940 | **<0.001** | -0.507 | -0.585 – -0.428 | **<0.001** | -11.44 | -12.95 – -9.93 | **<0.001** |
| timestep | 4.642 | 4.013 – 5.270 | **<0.001** | 0.223 | 0.200 – 0.247 | **<0.001** | 3.453 | 3.015 – 3.892 | **<0.001** |
| mother ID [2] | -0.082 | -0.216 – 0.051 | 0.227 | -0.012 | -0.025 – 0.000 | 0.057 | -0.164 | -0.329 – 0.001 | 0.051 |
| mother ID [3] | -0.012 | -0.081 – 0.058 | 0.739 | 0.002 | -0.005 – 0.008 | 0.655 | -0.001 | -0.088 – 0.085 | 0.979 |
| tank position [middle] | 0.027 | -0.053 – 0.108 | 0.504 | 0.001 | -0.007 – 0.008 | 0.887 | 0.032 | -0.068 – 0.133 | 0.529 |
| tank position [wall] | 0.022 | -0.063 – 0.107 | 0.614 | -0.003 | -0.011 – 0.005 | 0.465 | -0.055 | -0.161 – 0.052 | 0.314 |
| tank system [2] | 0.048 | -0.043 – 0.139 | 0.298 | 0.004 | -0.005 – 0.012 | 0.414 | 0.129 | 0.016 – 0.242 | **0.026** |
| tank system [3] | 0.029 | -0.067 – 0.126 | 0.551 | -0.000 | -0.010 – 0.009 | 0.926 | 0.012 | -0.108 – 0.132 | 0.842 |
| tank system [4] | -0.001 | -0.093 – 0.091 | 0.984 | -0.004 | -0.012 – 0.005 | 0.430 | 0.044 | -0.070 – 0.158 | 0.452 |
| **Random Effects** | | | | | | | | | |
| $\sigma^2$ | 0.04234 | | | 0.00005 | | | 0.02089 | | |
| $\tau_{00}$ | 0.00758 $_{id.2}$ | | | 0.00010 $_{id.2}$ | | | 0.01681 $_{id.2}$ | | |
| $\tau_{11}$ | 0.37576 $_{id.timestep}$ | | | 0.00180 $_{id.timestep}$ | | | 0.15382 $_{id.timestep}$ | | |
| | 6.68184 $_{id.I(timestep^2)}$ | | | 0.01845 $_{id.I(timestep^2)}$ | | | 3.12505 $_{id.I(timestep^2)}$ | | |
| ICC | 0.19557 | | | 0.50830 | | | 0.16787 | | |
| N | 45 $_{id}$ | | | 45 $_{id}$ | | | 45 $_{id}$ | | |
| Observations | 1239 | | | 1239 | | | 1239 | | |
| Marginal $R^2$ / Conditional $R^2$ | 0.151 / 0.317 | | | 0.399 / 0.705 | | | 0.281 / 0.402 | | |

**Supplementary Table 3.3.** Output for linear and quadratic models of behavioral entropy (calculated on a grid of 20 clusters obtained via UMAP dimension reduction and watershed segmentation) regressed against time (in days) as the continuous fixed predictor of interest. The quadratic model contains an additional term for time$^2$ as a continuous fixed predictor. Mother ID, tank position, and tank system are included as categorical fixed predictors, and fish ID is included as a random factor with varying intercepts and linear coefficients (as well as varying quadratic coefficients in the case of the quadratic model).

| | **Behavioral entropy (linear model – day interval)** | | | **Behavioral entropy (quadratic model – day interval)** | | |
|---|---|---|---|---|---|---|
| *Predictors* | *Estimates* | *CI* | *p* | *Estimates* | *CI* | *p* |
| (Intercept) | 1.79 | 1.65 – 1.92 | **<0.001** | 1.76 | 1.32 – 2.21 | **<0.001** |
| timestep | 0.87 | 0.21 – 1.53 | **0.010** | 7.16 | 5.45 – 8.88 | **<0.001** |
| mother ID [2] | -0.10 | -0.19 – -0.00 | **0.049** | -1.08 | -1.69 – -0.47 | **0.001** |
| mother ID [3] | -0.07 | -0.12 – -0.02 | **0.006** | -0.26 | -0.58 – 0.06 | 0.113 |
| tank position [middle] | 0.01 | -0.05 – 0.06 | 0.819 | -0.06 | -0.43 – 0.32 | 0.771 |
| tank position [wall] | 0.02 | -0.04 – 0.08 | 0.562 | -0.24 | -0.63 – 0.16 | 0.240 |
| tank system [2] | 0.06 | -0.00 – 0.13 | 0.064 | 0.26 | -0.16 – 0.68 | 0.225 |
| tank system [3] | 0.00 | -0.07 – 0.07 | 0.912 | -0.14 | -0.58 – 0.31 | 0.549 |
| tank system [4] | -0.03 | -0.10 – 0.04 | 0.368 | -0.15 | -0.58 – 0.27 | 0.475 |
| timestep^2 | | | | -21.67 | -26.08 – -17.25 | **<0.001** |
| **Random Effects** | | | | | | |
| $\sigma^2$ | 0.07 | | | 0.05 | | |
| $\tau_{00}$ | 0.16 $_{id}$ | | | 0.25 $_{id.2}$ | | |
| $\tau_{11}$ | 4.63 $_{id.timestep}$ | | | 29.14 $_{id.timestep}$ | | |
| | | | | 170.19 $_{id.I(timestep\wedge 2)}$ | | |
| ICC | 0.35 | | | 0.94 | | |
| N | 45 $_{id}$ | | | 45 $_{id}$ | | |
| Observations | 1239 | | | 1239 | | |
| Marginal $R^2$ / Conditional $R^2$ | 0.061 / 0.385 | | | 0.128 / 0.948 | | |

**Supplementary Table 3.4.** Table of Akaike Information Criteria (AIC) and Bayesian Information Criteria (BIC) used to compare linear and quadratic models using day-length intervals (see Supplemental Tables 3.1 and 3.2) for each response.

| **Model** | **ΔAIC** | **ΔBIC** |
|---|---|---|
| Step length CoV quadratic (day interval) | 0.00 | 0.00 |
| Step length CoV linear (day interval) | 163.05 | 157.93 |
| Turning angle CoV quadratic (day interval) | 0.00 | 0.00 |
| Turning angle CoV linear (day interval) | 149.24 | 144.12 |
| Distance to tank wall CoV quadratic (day interval) | 0.00 | 0.00 |
| Distance to tank wall CoV linear (day interval) | 201.18 | 196.06 |
| Behavioral entropy quadratic (day interval) | 0.00 | 0.00 |
| Behavioral entropy linear (day interval) | 172.00 | 166.87 |

**Supplementary Figure 3.1**. Developmental arcs of behavioral plasticity calculated as either the coefficient of variation in a single behavioral metric (A-C) or as a measure of behavioral entropy across the entire movement repertoire (D) for the first 28 days of development. In contrast to Fig 1 and Fig 3 in the main text, which calculates coefficients of variation or entropy over one-hour intervals, this figure corresponds to coefficients of variation or entropy that were calculated over one-day intervals. Note that, over these scales, the length of time over which data were aggregated has no bearing on the overall qualitative results (see also Supplemental Tables 3.1-4).

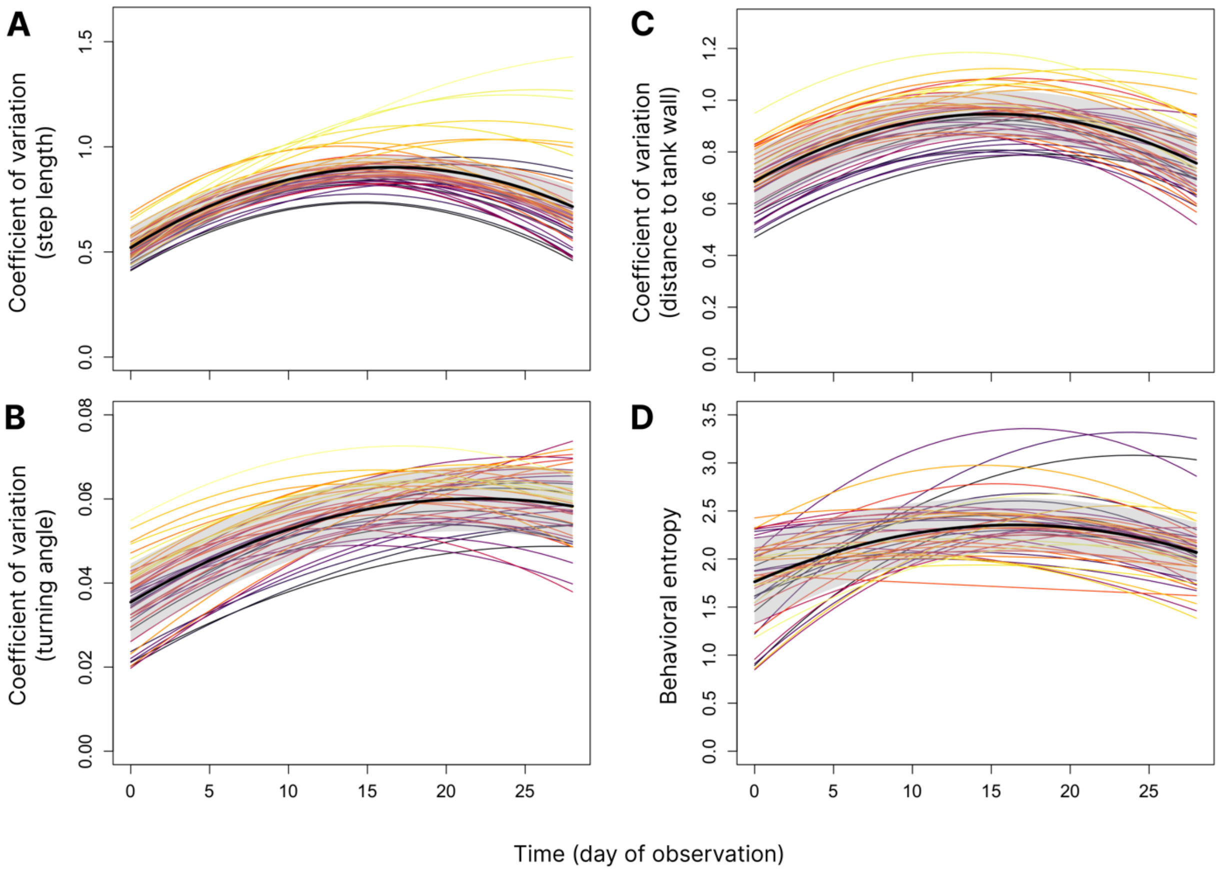

#### Supplement 4: Behavioral entropy measures calculated across varying cluster numbers and with multiple clustering algorithms

**Supplementary Table 4.1.** Output for linear models of behavioral entropy (calculated on a grid of 7 and 10 clusters obtained via UMAP dimension reduction and watershed segmentation) regressed against time (in hours) as the continuous fixed predictor of interest. Mother ID, tank position, and tank system are included as categorical fixed predictors, and fish ID is included as a random factor with varying intercepts and slopes.

| | **Behavioral entropy – 7 clusters (linear model)** | | | **Behavioral entropy – 10 clusters (linear model)** | | |
|---|---|---|---|---|---|---|
| *Predictors* | *Estimates* | *CI* | *p* | *Estimates* | *CI* | *p* |
| (Intercept) | 0.61 | 0.53 – 0.69 | **<0.001** | 1.36 | 1.25 – 1.46 | **<0.001** |
| timestep | 0.03 | -0.00 – 0.06 | 0.081 | 0.06 | -0.00 – 0.12 | 0.070 |
| mother ID [2] | -0.01 | -0.12 – 0.09 | 0.800 | -0.10 | -0.19 – -0.01 | **0.024** |
| mother ID [3] | -0.20 | -0.26 – -0.15 | **<0.001** | -0.12 | -0.17 – -0.08 | **<0.001** |
| tank position [middle] | -0.01 | -0.07 – 0.05 | 0.724 | -0.00 | -0.06 – 0.05 | 0.954 |
| tank position [wall] | 0.01 | -0.05 – 0.08 | 0.716 | 0.00 | -0.05 – 0.06 | 0.873 |
| tank system [2] | 0.02 | -0.05 – 0.09 | 0.661 | 0.06 | 0.00 – 0.13 | **0.041** |
| tank system [3] | -0.01 | -0.08 – 0.07 | 0.812 | 0.01 | -0.06 – 0.07 | 0.844 |
| tank system [4] | -0.01 | -0.08 – 0.06 | 0.813 | -0.02 | -0.08 – 0.04 | 0.472 |
| **Random Effects** | | | | | | |
| $\sigma^2$ | 0.06 | | | 0.09 | | |
| $\tau_{00}$ | 0.02 $_{id}$ | | | 0.09 $_{id}$ | | |
| $\tau_{11}$ | 0.01 $_{id.timestep}$ | | | 0.05 $_{id.timestep}$ | | |
| ICC | 0.17 | | | 0.23 | | |
| N | 45 $_{id}$ | | | 45 $_{id}$ | | |
| Observations | 9861 | | | 9861 | | |
| Marginal $R^2$ / Conditional $R^2$ | 0.129 / 0.280 | | | 0.051 / 0.273 | | |

**Supplementary Table 4.2.** Output for quadratic models of behavioral entropy (calculated on a grid of 7 and 10 clusters obtained via UMAP dimension reduction and watershed segmentation) regressed against time (in hours) as the continuous fixed predictor of interest, including a time$^2$ quadratic term. Mother ID, tank position, and tank system are included as categorical fixed predictors, and fish ID is included as a random factor with varying intercepts, linear, and quadratic coefficients.

| | **Behavioral entropy – 7 clusters (quadratic model)** | | | **Behavioral entropy – 10 clusters (quadratic model)** | | |
|---|---|---|---|---|---|---|
| *Predictors* | *Estimates* | *CI* | *p* | *Estimates* | *CI* | *p* |
| (Intercept) | 0.50 | 0.39 – 0.61 | **<0.001** | 1.38 | 1.02 – 1.74 | **<0.001** |
| timestep^2 | -0.11 | -0.15 – -0.07 | **<0.001** | -0.27 | -0.34 – -0.20 | **<0.001** |
| timestep | 0.29 | 0.20 – 0.37 | **<0.001** | 0.66 | 0.47 – 0.86 | **<0.001** |
| mother ID [2] | -0.17 | -0.32 – -0.02 | **0.030** | -0.87 | -1.37 – -0.38 | **0.001** |
| mother ID [3] | -0.10 | -0.18 – -0.02 | **0.014** | -0.23 | -0.48 – 0.03 | 0.088 |
| tank position [middle] | 0.00 | -0.09 – 0.10 | 0.919 | -0.08 | -0.39 – 0.22 | 0.580 |
| tank position [wall] | 0.03 | -0.06 – 0.13 | 0.488 | -0.24 | -0.56 – 0.08 | 0.138 |
| tank system [2] | -0.04 | -0.14 – 0.06 | 0.460 | 0.21 | -0.13 – 0.54 | 0.234 |
| tank system [3] | -0.09 | -0.20 – 0.02 | 0.115 | -0.15 | -0.51 – 0.21 | 0.413 |
| tank system [4] | -0.03 | -0.13 – 0.08 | 0.612 | -0.15 | -0.49 – 0.19 | 0.400 |
| **Random Effects** | | | | | | |
| $\sigma^2$ | 0.06 | | | 0.07 | | |
| $\tau_{00}$ | $0.01_{id.2}$ | | | $0.17_{id.2}$ | | |
| $\tau_{11}$ | $0.08_{id.timestep}$ | | | $0.44_{id.timestep}$ | | |
| | $0.02_{id.I(timestep\text{^}2)}$ | | | $0.05_{id.I(timestep\text{^}2)}$ | | |
| ICC | 0.70 | | | 0.91 | | |
| N | $45_{id}$ | | | $45_{id}$ | | |
| Observations | 9861 | | | 9861 | | |
| Marginal $R^2$ / Conditional $R^2$ | 0.037 / 0.708 | | | 0.094 / 0.920 | | |

**Supplementary Table 4.3.** Output for linear and quadratic models of behavioral entropy (calculated on a grid of 20 clusters obtained via k-means clustering) regressed against time (in hours) as the continuous fixed predictor of interest. The quadratic model contains an additional quadratic term for time$^2$ as a continuous fixed predictor. Mother ID, tank position, and tank system are included as categorical fixed predictors, and fish ID is included as a random factor with varying intercepts and linear coefficients (as well as varying quadratic coefficients in the case of the quadratic model).

| | **Behavioral entropy calculated with k-means (linear model)** | | | **Behavioral entropy calculated with k-means (quadratic model)** | | |
|---|---|---|---|---|---|---|
| *Predictors* | *Estimates* | *CI* | *p* | *Estimates* | *CI* | *p* |
| (Intercept) | 1.91 | 1.79 – 2.04 | **<0.001** | 1.94 | 1.52 – 2.37 | **<0.001** |
| timestep | 0.07 | 0.00 – 0.15 | **0.038** | 0.80 | 0.57 – 1.02 | **<0.001** |
| mother ID [2] | -0.08 | -0.17 – 0.02 | 0.114 | -1.07 | -1.66 – -0.49 | **<0.001** |
| mother ID [3] | -0.13 | -0.18 – -0.08 | **<0.001** | -0.25 | -0.56 – 0.06 | 0.115 |
| tank position [middle] | 0.01 | -0.05 – 0.07 | 0.684 | -0.11 | -0.47 – 0.25 | 0.543 |
| tank position [wall] | 0.02 | -0.04 – 0.08 | 0.488 | -0.27 | -0.65 – 0.11 | 0.166 |
| tank system [2] | 0.04 | -0.02 – 0.11 | 0.200 | 0.24 | -0.16 – 0.65 | 0.243 |
| tank system [3] | 0.01 | -0.06 – 0.08 | 0.721 | -0.18 | -0.61 – 0.24 | 0.401 |
| tank system [4] | -0.02 | -0.08 – 0.05 | 0.659 | -0.16 | -0.57 – 0.24 | 0.432 |
| timestep^2 | | | | -0.32 | -0.40 – -0.24 | **<0.001** |
| **Random Effects** | | | | | | |
| $\sigma^2$ | 0.11 | | | 0.09 | | |
| $\tau_{00}$ | 0.13 $_{id}$ | | | 0.24 $_{id.2}$ | | |
| $\tau_{11}$ | 0.06 $_{id.timestep}$ | | | 0.59 $_{id.timestep}$ | | |
| | | | | 0.07 $_{id.I(timestep^2)}$ | | |
| ICC | 0.25 | | | 0.92 | | |
| N | 45 $_{id}$ | | | 45 $_{id}$ | | |
| Observations | 9861 | | | 9861 | | |
| Marginal $R^2$ / Conditional $R^2$ | 0.046 / 0.285 | | | 0.099 / 0.927 | | |

**Supplementary Figure 4.1**. Developmental arcs of plasticity in movement repertoires of Amazon mollies over the first 28 days of life, quantified as behavioral entropy. Shown here are graphs of behavioral entropy through time, where entropy is calculated using a range of cluster numbers and over two alternative clustering algorithms. Note that the general pattern of entropy remains unchanged across cluster numbers and specific clustering algorithms: (A) entropy calculated across only 7 clusters, obtained via UMAP dimension reduction and watershed segmentation, (B) entropy calculated across 10 clusters, obtained via UMAP dimension reduction and watershed segmentation, (C) entropy calculated across 20 clusters, obtained via UMAP dimension reduction and watershed segmentation (corresponding to Figure 3 in the main text), and (D) entropy calculated across 20 clusters, obtained via k-means clustering.

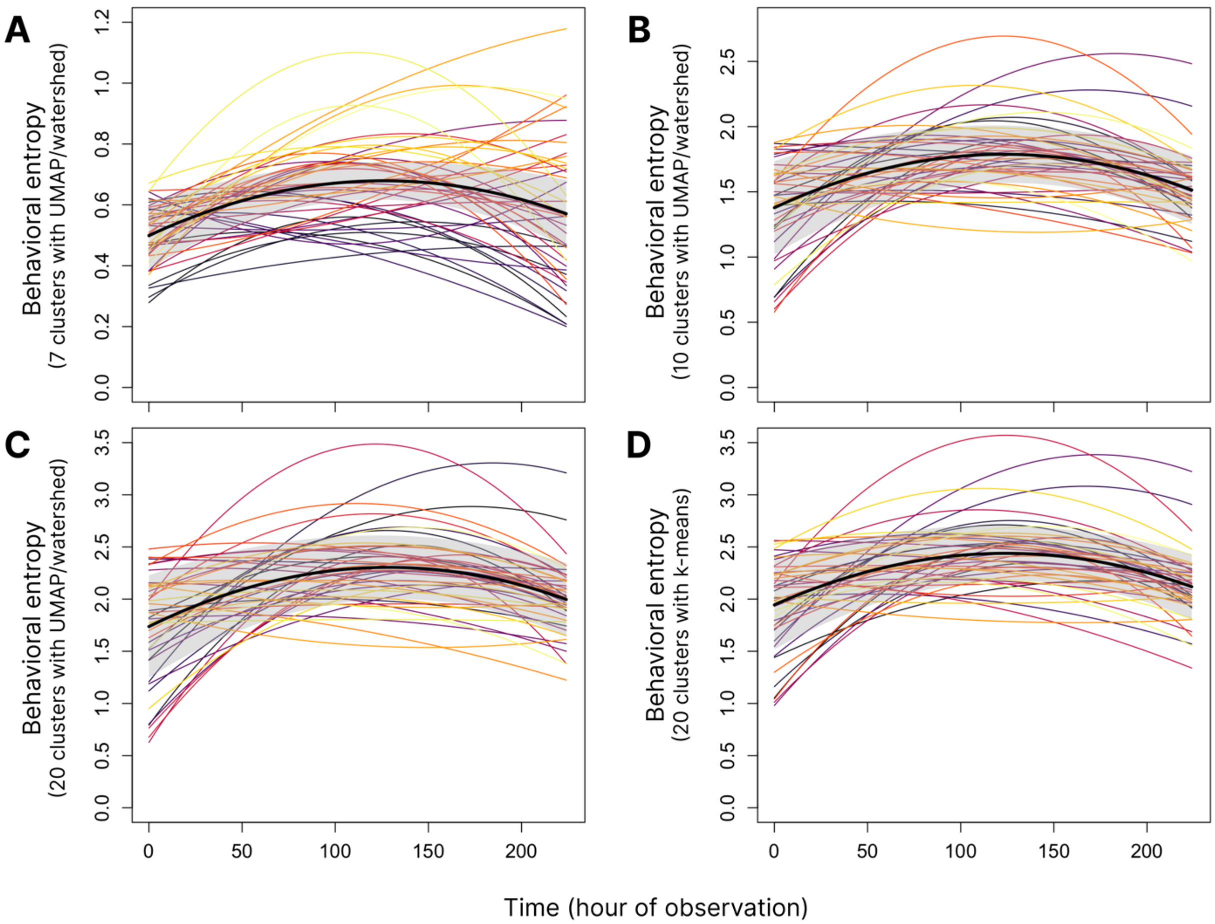

#### Supplement 5: Movie of simultaneous movement in physical space and behavioral phenotype space for low- and high-entropy individuals

**Supplementary Movie 5.1.** A 5-minute movie in which three low entropy individuals (A) and three high entropy individuals (B) are shown exhibiting a continuous track of behavior over a 30-minute real-time sequence (6x speed) in both physical tank space (triangular shapes on the left of each panel) and clustered behavioral phenotype (UMAP) space. Individuals are color-coded.
